# Bile acid-mediated ileal barrier function deficits in primary sclerosing cholangitis

**DOI:** 10.64898/2026.11.25.768979

**Authors:** Miriam Tomczak, Katherine Coldwell, Kyla Dooley, Amelia Heslington, Claire Pearson, Elizete Araujo, Megan Perry, Kaye Eve-Rowe, Shana-Lee Bownes, Yukari Kimura, Nicholas McDermott Provine, Lisa Gartner, Amy Cross, Joanna Hester, Iolanda Vendrell, Anna Hoyle, Roman Fischer, Sandi Yen, Charlotte Rich-Griffin, Ida Parisi, Oyindamola Ajisegiri, Ethar Ali, Sumaiya Hoque, TGLU investigators, Human Cell Atlas (HCA) investigators, Fadi Issa, Samuel Huber, Fiona Powrie, Matthias Friedrich, Holm Hans Uhlig, Calliope Dendrou, Emma Louise Culver, Paul Klenerman, Nicholas Ilott

## Abstract

**Background:** Given a high prevalence of ulcerative colitis (UC) in primary sclerosing cholangitis (PSC-UC), research into the pathophysiology of the gut-liver axis has focussed on the large intestine. However, reduced bile flow from the liver has the potential to impact on homeostatic functions of the small intestine, including the main site of bile acid reuptake – the terminal ileum.

**Objective:** Define ileal barrier changes in PSC-UC that are future treatment targets.

**Design:** We used a combination of single-cell RNA-sequencing, spatial transcriptomics, bulk RNA-seq, and liquid chromatography mass spectrometry (LC-MS)-based bulk proteomics in samples from patients with PSC-UC, UC and healthy controls to determine PSC-UC-associated barrier function changes in the terminal ileum.

**Results:** Mature enterocytes of the terminal ileum are characterised by spatially organised cell states. The phase II detoxification gene, *Glutathione-S-transferase alpha 1* (*GSTA1*), marks one state that is enriched at the base of villi and is co-expressed with detoxification genes. Genes within this co-expression module are downregulated in ileal samples from patients with PSC-UC and protein abundance is correlated with liver function in patients. We confirm that *Gsta1* mRNA is decreased in the ileum in murine cholestatic liver disease and altering microbiome capacity for bile acid deconjugation regulates *Gsta1* expression.

**Conclusion:** We have identified new pathways that are disrupted in the ileum of patients with PSC-UC which represent potential new targets for microbiome-based treatments to improve barrier function.

## Introduction

Primary sclerosing cholangitis (PSC) is a rare progressive disease of the bile ducts and liver with no effective medical treatments. The impacts of restricted bile flow (cholestasis) in PSC are wide-ranging. Retention of toxic concentrations of bile acids in the liver results in direct liver damage, where decreased concentrations reaching target organs such as the small and large intestine impact microbial ecology, nutrient absorption, intestinal barrier function and immunity. Combined with the major impact of ulcerative colitis (UC) in patients with PSC,^1^ these features support a role for gut-liver communication in disease progression.

Causal relationships between the liver and gut in mediating pathology in PSC remain unclear. Studies in mice have revealed both pathogenic and protective effects of gut-derived factors and inflammation on liver pathology,^2,3^ and it is likely that cholestasis itself impacts downstream functions of the intestine that may contribute to – or protect against – gut inflammation.^4^ Recent focus of research has been the impact of the gut microbiome on liver diseases that include PSC and PSC -UC.^5–22^ Coupling of increased intestinal permeability and pathobiont expansion may contribute to disease progression in PSC. Indeed, murine studies have highlighted a pathogenic role for the microbial members *Klebsiella pneumoniae* and *Enterococcus spp.* in driving progression of liver disease through bacterial translocation,^23^ as well as the pathogenic impact of LPS-driven macrophage-derived Il1b on liver fibrosis.^3^ Moreover, bacteria-derived outer-membrane vesicles can translocate from the gut to the liver, and may impact on liver function and pathology.^16^ However, the exact mechanisms through which bacteria in the gut are capable of imparting long-range impact on the liver remain incompletely understood.

Tight control of the intestinal barrier through constant replenishment of mature epithelial cells from stem cell progenitors, as well as homeostatic innate and adaptive immune responses to commensal bacteria prevent pathogenic translocation and risk of systemic responses to gut bacterial products.^24^ However, disruption to barrier function through disease processes such as chronic inflammation in IBD can lead to aberrant local and systemic immune responses to translocating commensal microbes.^24–27^ There is increasing recognition of the role of increased intestinal permeability and gut microbiota-derived factors in chronic liver diseases,^28^ which includes a protective role for conjugated bile acids on epithelial barrier integrity.^29^ The presence of large bowel inflammation in the majority of patients with PSC has led to an overarching hypothesis that pathophysiologic gut-liver communication and increased intestinal permeability are the result of a breakdown of the colonic barrier in PSC. While this route of communication may lead to inflammation-driven homing of memory T-cells from the colon to the liver through inflammation-driven upregulation of adhesion molecules such as vascular adhesion protein 1 (VAP-1),^30^ or the translocation of bacteria and their products, a role for the small intestine in PSC-associated liver disease has not been explored.

Given a fundamental role of the terminal ileum in controlling bile acid biology and the potential for reduced bile flow to impact ileal barrier function in PSC-UC, we aimed to explore alterations to the transcriptome and proteome in patients with PSC-UC relative to healthy controls and patients with UC. We describe key changes to detoxification programmes in ileal enterocytes in PSC-UC and confirm bile acid control of these processes, that can be tuned through microbial modulation of bile acid deconjugation.

## Results

### Ileal enterocytes have diverse cell states that are marked by high *GSTA1* or *APOA1* expression

To understand barrier-related changes in the terminal ileum of patient with PSC-UC, we generated multiple high-dimensional datasets across different patient cohorts (**Fig. 1A**). First, we analysed a single cell gene expression atlas (10X Chromium) of unsorted cells derived from terminal ileum biopsies from 4 patients with PSC-UC, 4 patients with UC alone and 4 healthy controls (**Table 1** and **Fig. S1A**). Using canonical marker genes and cluster marker genes (**Fig. S1B**), we found the major cell types identified were enterocytes (15.6%), CD4 T cells (14.60%), B cells (13.1%), CD8 T cells (8.5%), IgA-producing B cells (8.2%), monocytes/macrophages (5.8%) and epithelial stem cells (5.0%) that together made up 70.8% of cells (**Fig. S1C**). We used these broad classifications to subset and sub-cluster individual compartments, applying cell-type-specific clustering parameters and annotation to identify epithelial (**Fig. 1B**, n = 3,233), immune (**Fig. 1C**, n = 8,388) and stromal (**Fig. 1D**, n = 712) cells. The median read count per cell across samples ranged from 1,784 – 8,732.

**Fig. 1.**
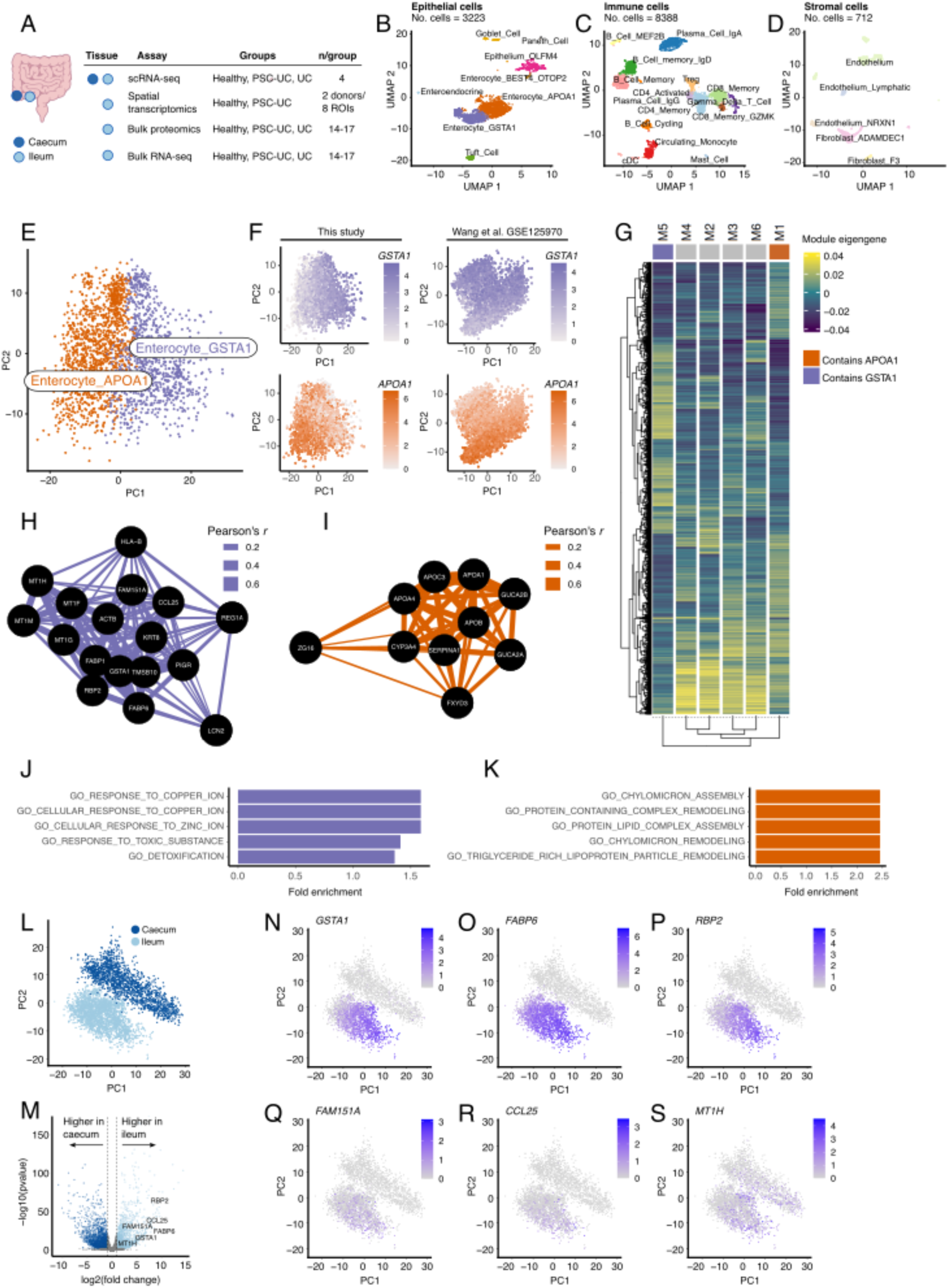
Ileal enterocytes are represented by two cell states. (A) Overview of human cohort studies presented in this manuscript. UMAP projections of annotated (B) epithelial cells (C) immune cells, and (D) stromal cells. (E-F) Principal component plots from our study and data from Wang *et al.* showing conserved cell state phenotypes. (G) Modules derived from scWGCNA analysis represented in a heatmap of module eigengenes. Networks of genes co-expressed with (H) *GSTA1* and (I) *APOA1* across ileal enterocytes. (J) GO:BP pathway enrichment in genes co-expressed with *GSTA1*. (K) GO:BP pathway enrichment in genes co-expressed with APOA1. (L) PCA plot showing mature absorptive epithelial cells from the ileum and the caecum. (M) Volcano plot showing the results of differential expression analysis (DESeq2) between enterocytes of the ileum vs. colonocytes of the caecum. Highlighted are those genes that are part of the *GSTA1* co-expression module. Tissue-specific expression of (N) *GSTA1*, (O) *FABP6*, (P) *RBP2*, (Q) *FAM151A*, (R) *CCL25* and (S) *MT1H*.

**Table 1.**
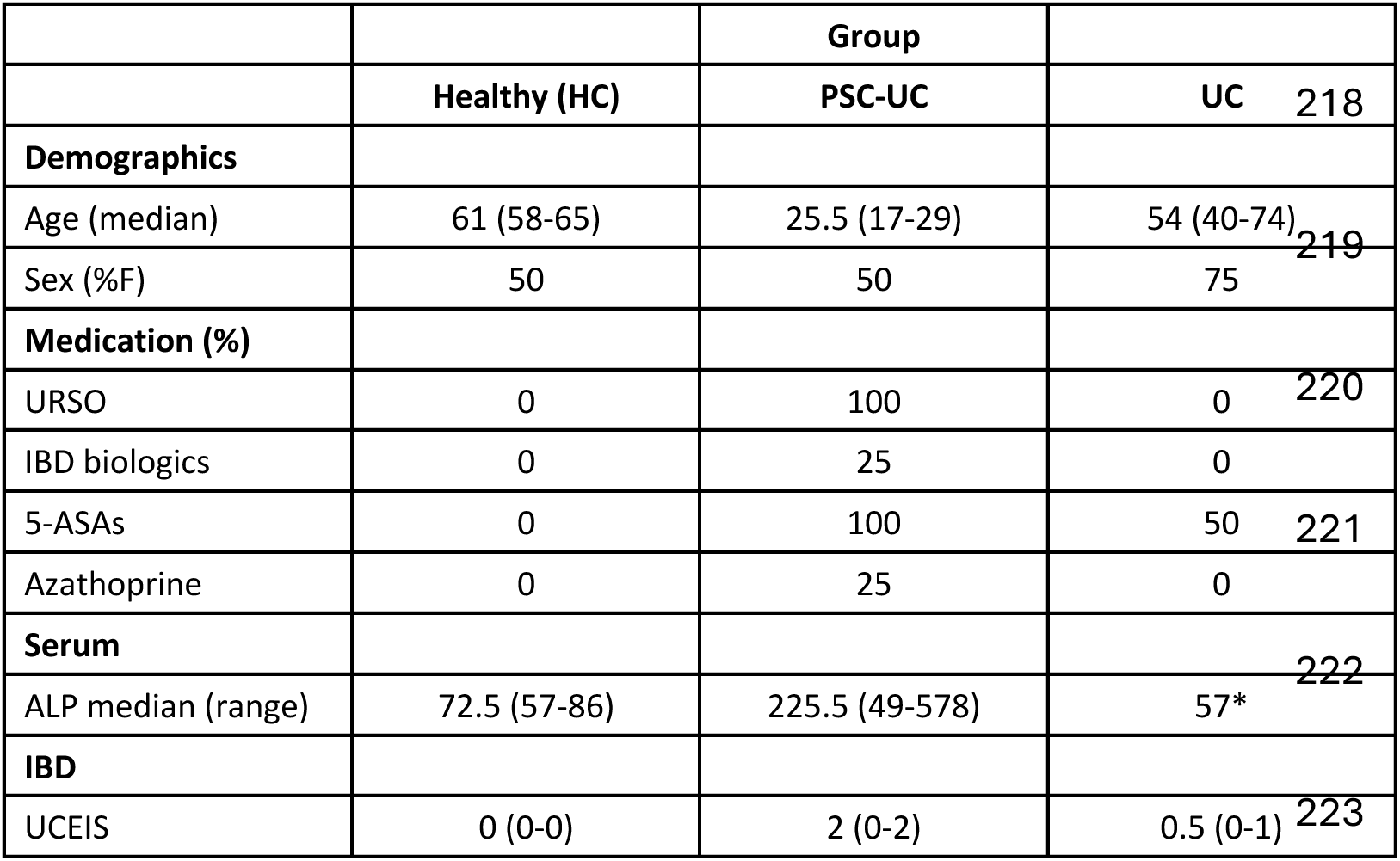
Characteristics of participants in the study (n = 4/group). *denotes that data were not available for 3/4 of the participants.

|  |  | **Group** |  |
|---|---|---|---|
|  | **Healthy (HC)** | **PSC-UC** | **UC** |
| **Demographics** |  |  |  |
| Age (median) | 61 (58-65) | 25.5 (17-29) | 54 (40-74) |
| Sex (%F) | 50 | 50 | 75 |
| **Medication (%)** |  |  |  |
| URSO | 0 | 100 | 0 |
| IBD biologics | 0 | 25 | 0 |
| 5-ASAs | 0 | 100 | 50 |
| Azathoprine | 0 | 25 | 0 |
| **Serum** |  |  |  |
| ALP median (range) | 72.5 (57-86) | 225.5 (49-578) | 57* |
| **IBD** |  |  |  |
| UCEIS | 0 (0-0) | 2 (0-2) | 0.5 (0-1) |

Given a central role of ileal epithelial cells in integrating environmental and immune signals (including bile acids from the liver), we focused our attention on this cell compartment. The majority of epithelial cells identified in our initial clustering were mature enterocytes.

Clustering and marker gene analysis revealed two subsets of mature enterocytes that were characterised by either high *GSTA1* or high *APOA1* expression (**Fig. 1B**), a feature of human terminal ileum enterocytes that has not been previously described. Sub-clustering of enterocyte subsets revealed that these cell types are overlapping and likely represent variant cell states (**Fig. 1E**). To determine if this is a common feature of human enterocytes, we utilised a publicly available scRNA-seq data set from healthy human ileum to assess their presence. In both our data set and that of Wang *et al.*^31^, we demonstrated a spectrum of *GSTA1* and *APOA1* expression across enterocytes, with the highest *GSTA1* expressing cells opposing the cells with the highest *APOA1* expression in a PCA projection (**Fig. 1F**). To further explore coordinated regulation and functional profiles of these cell states, we performed single-cell Weighted Gene Co-expression Network Analysis (scWGCNA). We detected that modules containing *GSTA1* (module 5) and *APOA1* (module 1) were opposed in their module eigengene expression across enterocytes, as would be expected (**Fig. 1G**).

Taking the most highly connected genes in each of these modules (module membership > 0.6), we demonstrated that genes correlated with *GSTA1* expression (**Fig. 1H**) were enriched for gene ontology (GO) biological pathways that represent metal-binding and detoxification (**Fig. 1J**), where genes that are most highly correlated with *APOA1* expression (**Fig. 1I**) are enriched for GO pathways related to chylomicron assembly (**Fig. 1K**).

Genes co-expressed with *GSTA1* included metallothioneins (*MT1F*, *MT1G*, *MT1H* and *MT1M*), that code for cysteine-rich proteins capable of binding heavy metals and preventing heavy metal toxicity, as well as acting as reducing agents for the detoxification of reactive oxygen species.^32^ Of note, *GSTA1*-associated genes also included genes involved in bile acid (BA) binding and transport (*FABP6, FABP1*), supporting a potential link between BA signalling and cellular detoxification pathways. Genes co-expressed with *APOA1* included multiple additional apolipoproteins (*APOA4, APOC3* and *APOB*) that are major players in transporting dietary lipids from the ileum through formation of high- and low-density lipoproteins and chylomicrons.

We additionally profiled 1,960 epithelial cells from the caecum in our cohort and explored the presence or absence of *GSTA1-* and *APOA1*-associated cell states in caecal colonocytes.

Mature absorptive epithelial cells (colonocytes in the caecum and enterocytes in the ileum) clustered separately in PCA (**Fig. 1L**), corroborating functional specialisation of these cells at each tissue location. Of note, genes associated with *GSTA1*, including *GSTA1* itself, are expressed at lower levels in caecal colonocytes than ileal enterocytes (**Fig. 1M**), and display largely tissue-restricted expression patterns (**Fig. 1N-S**). This suggests that *GSTA1-*associated gene expression programmes are specific to small intestinal function, which may relate to BA signalling.

### Ileal enterocyte cell states are spatially partitioned

We wanted to further explore the relationship between our defined enterocyte cell states and the intestinal niche. Given previous analysis in enterocytes of the human proximal small intestine that supported zonation of *GSTA1* and chylomicron assembly genes along the crypt- villus axis,^33^ we used the Nanostring GeoMx digital spatial profiler (DSP) platform to assess the spatial arrangement of *GSTA1-* and *APOA1*-associated gene modules to determine partitioning in the ileum. Using DSP segmentation and high-throughput sequencing, we acquired whole transcriptomic data from PanCK+ cells in these regions (**Fig. 2A**). These data provided transcriptional information on 47% of *GSTA1*-associated genes (Module 5) and 64% of *APOA1*-associated genes (Module 1). We filtered the gene expression matrix for these genes and showed that we could separate base and tip regions of interest (ROI) based on this profile (**Fig. 2B**, Adonis test *R*^2^ = 0.38, *p* = 0.03). Using linear mixed-effects models (with ROI type as the fixed effect and donor as the random effect), we observed nominal significance for a number of genes in these modules and their association with either the base or the tip of villi (**Fig. 2C**, including *GSTA1* and *APOA1*). Visualisation of these gene sets revealed an association between *GSTA1*-associated genes with the base of villi and *APOA1*-associated genes with the villus tip (**Fig. 2C**). These data were further confirmed using publicly available Visium data on an human ileum biopsy, where we observed expression of *APOA1* at villus tips and *GSTA1* at the villus base (**Fig. 2D**).

**Fig. 2.**
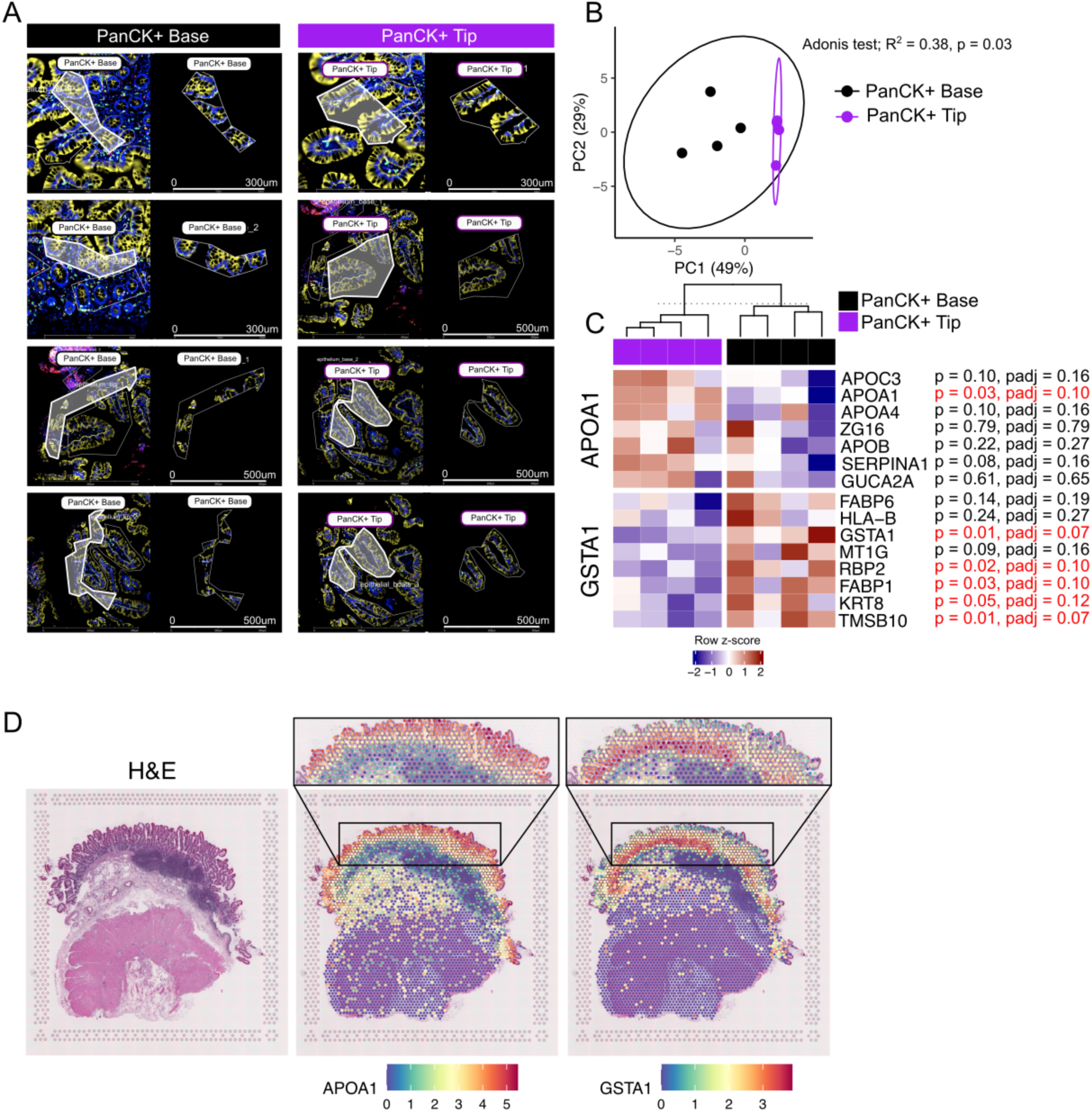
Enterocyte cell states are spatially organised in the ileum. (A) Regions of interest defined on the Nanostring Digital Spatial Profiler (DSP). Regions were draw free-hand around the villus tips and bases based guided by PanCK+ staining and crypt structure. (B) Principal component plot based on expression of *APOA1* and *GSTA1* co-expressed genes. (C) Heatmap showing normalised expression values for genes that are part of the *APOA1* or *GSTA1* co-expressed gene modules annotated by expression in PanCK+ cells at the tip or base of villi. (D) Expression of *APOA1* and *GSTA1* in a publicly available Visium dataset. Values are normalised values from Seurat V5. Note that zoomed in regions approximate the locations in the full image.

These data extend previous observations in the proximal small intestine with evidence of partitioning of human ileal enterocyte cell states into base-associated detoxification and villus-associated lipid transport states.

### Enterocyte *GSTA1*-associated detoxification pathways are decreased in PSC-UC

We next sought to define the genes and pathways that are dysregulated in the epithelial compartment in PSC-UC. We did not identify any statistically significant changes in proportions of epithelial cell types in PSC-UC. Two samples (n = 1 PSC-UC and n = 1 UC) displayed high proportions of epithelial cells that were annotated as *OLFM4*-expressing epithelial cells (**Fig. 3A**) and given a lack of representative enterocyte cell types in these samples (**Fig. S2**), they were not included in downstream analysis.

**Fig. 3.**
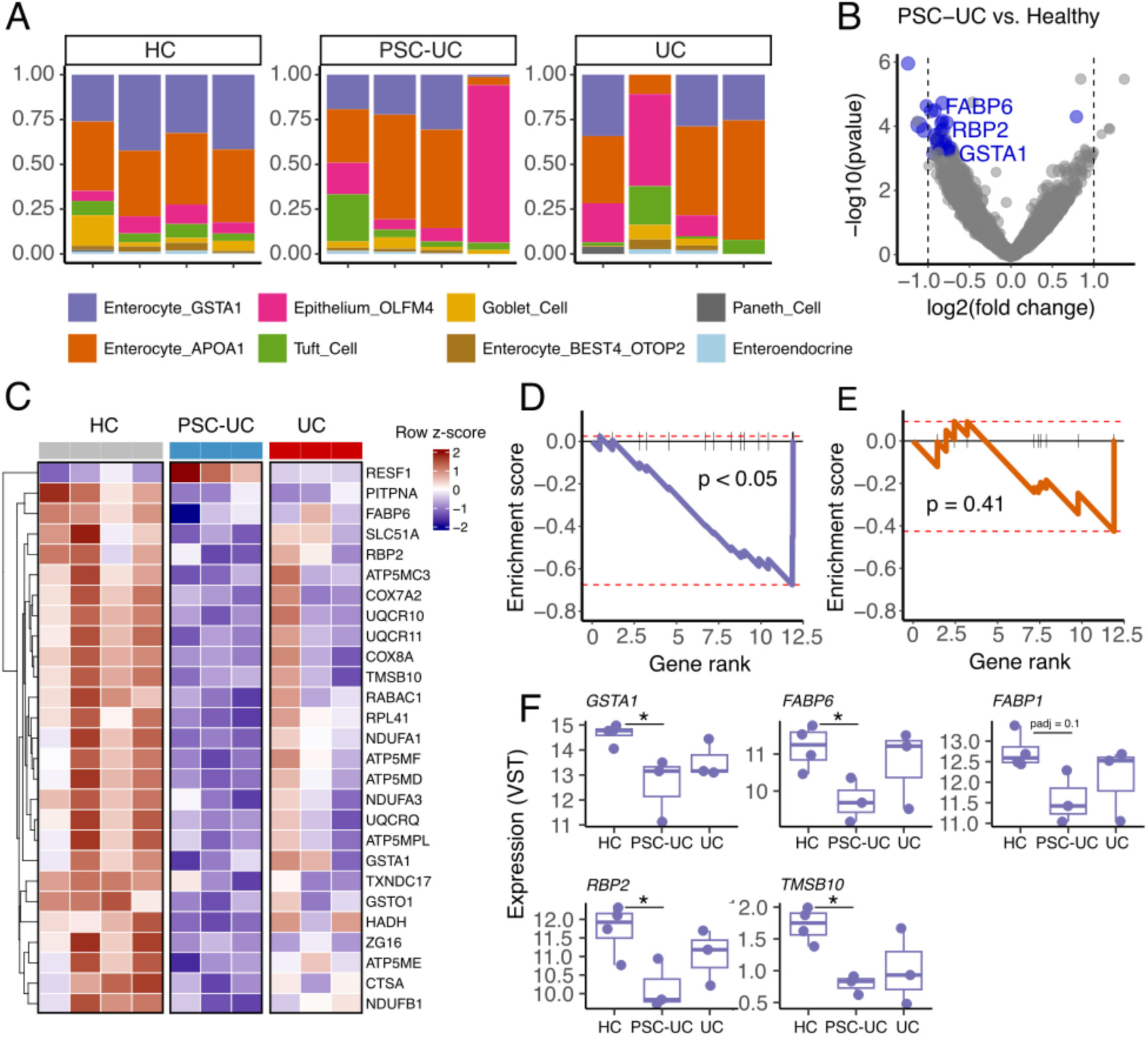
Genes enriched for detoxification pathways are decreased in the ileum of patients with PSC-UC. (A) Proportion of each epithelial cell type identified in each sample and across disease groups. (B) Volcano plot displaying differential expression results from pseudobulk analysis in enterocytes. (C) Heatmap of normalised (VST and row-scaled) expression of genes called as differentially expressed in PSC-UC relative to healthy controls. (D) Gene set enrichment analysis of *GSTA1* co-expressed genes against genes ranked by fold change in PSC-UC vs Healthy controls. (E) Gene set enrichment analysis of *APOA1* co-expressed genes against genes ranked by fold change in PSC-UC vs Healthy controls. (F) Gene expression across disease groups of genes that form the leading-edge set of genes in the *GSTA1* module from (D).

To identify enterocyte gene expression associations in PSC-UC, we performed pseudobulk differential expression analysis on the two major enterocyte subsets (Enterocyte_APOA1 and Enterocyte_GSTA1) pooled together. We found 27 genes significantly decreased in PSC-UC relative to healthy controls and 1 increased (**Fig. 3B**). Fewer genes were identified in UC relative to healthy controls, with 1 decreased and 2 increased. Of note, 5/27 (36%) of genes that were identified as decreased in PSC-UC were members of our previously defined *GSTA1*-associated co-expression module (**Fig. 3C** *GSTA1, FABP6, RBP2, FABP1* and *TMSB10*) and implicated altered bile acid signalling (*FABP6, FABP1*), detoxification (*GSTA1*) and epithelial cell differentiation (*RBP2*) in PSC-UC. GSEA also revealed significant enrichment of *GSTA1*-associated genes amongst genes downregulated in PSC-UC relative to healthy controls (**Fig. 3D**), which was not the case for *APOA1*-associated genes.

Leading-edge genes (i.e. those contributing to the enrichment signal) reflected this module’s enrichment of detoxification pathways (*MT1M* and *GSTA1*) and its links to bile acid homeostasis (*FABP1* and *FABP6*). As expected, due to reduced bile flow in PSC-UC relative to healthy controls, we also identified decreased expression of additional bile acid-regulated genes in PSC-UC that include the canonical bile acid transporter *SLC10A2*, two bile-acid binding and export proteins, *SLC51A* and *SLC51B*, as well as the downstream enterohepatic bile acid feedback gene *FGF19*, at nominal p-value < 0.05 (**Fig. S3A-D**). A single gene, *ADIRF*, was significantly downregulated in PSC-UC caecum, which was similarly downregulated in ileum (**Fig. S4A**).

Together, these data support dysregulated detoxification pathways in enterocytes of patients with PSC-UC that are potentially linked to altered bile acid homeostasis.

### GSTA1 and associated detoxification proteins are linked to liver function in PSC-UC

We next investigated the regulation of *GSTA1* and its associated genes in the ileum of patients with PSC-UC and how this relates to protein abundance. We collected ileal biopsy samples from an additional cohort of patients diagnosed with PSC-UC (n = 14), UC only (n = 17) or healthy controls (n = 14) and performed combined bulk RNA-seq and LC-MS based proteomics (**Fig. 1A** and **Table 2**).

**Table 2.** Characteristics of participants in the study (n = 14-17/group).

|  | Group |  |  |
|---|---|---|---|
|  | **Healthy (HC) (n = 14)** | **PSC-UC (n = 14)** | **UC (n = 17)** |
| **Demographics** |  |  |  |
| Age median (SD) | 55 (15.21) | 37 (22.10) | 40 (18.16) |
| Sex (%F) | 57 | 43 | 47 |
| **Medication (%)** |  |  |  |
| URSO | 0 | 36 | 0 |
| IBD biologics | 0 | 0 | 18 |
| 5-ASAs | 0 | 29 | 41 |
| Azathoprine | 0 | 7 | 6 |
| **Serum** |  |  |  |
| ALP median (SD) | 74 (19.06) | 120 (196.92) | 75.5 (25.93) |

In a comparison of 6080 genes/proteins that were shared between RNA-seq and proteomics datasets, we observed a moderate and significant correlation between mean RNA expression and mean protein abundance across features (**Fig. 4A**). Of note, and excepting KTR8, mRNA expression of genes of interest in enterocytes (*GSTA1-*associated module genes) were correlated with protein levels across individuals (**Fig. 4B**), supporting that protein levels are tightly coupled to transcription for these genes. In this cohort, we observed significant variation in alkaline phosphatase levels in patients with PSC-UC (**Fig. S5A**). Thus, for downstream analysis, we performed statistical tests in two ways: 1) keeping all patients with PSC-UC as a single group and 2) splitting patients with PSC-UC into two groups (ALP High (>1.5x upper limit of the healthy control ALP U/L) and ALP Low (<1.5x upper limit of the healthy control ALP U/L)). We compared both protein and mRNA levels between groups, including sex and age as covariates. Of note, when considering patients with PSC-UC as a single group, we did not observe statistically significant differences in any genes/proteins that were part of the *GSTA1*-co-expressed module. Further, we did not observe any global shifts in the ileal proteome and transcriptome between diagnoses, irrespective of ALP classification (**Fig. 4C** and **D**). However, levels of GSTA1, MT1F and MT1H protein, and *GSTA1*, *MT1F* and *MT1H* mRNA were reduced in patients with high ALP levels (**Fig. 4E** and **F**, nominal p < 0.05) compared to healthy controls.

**Fig. 4.**
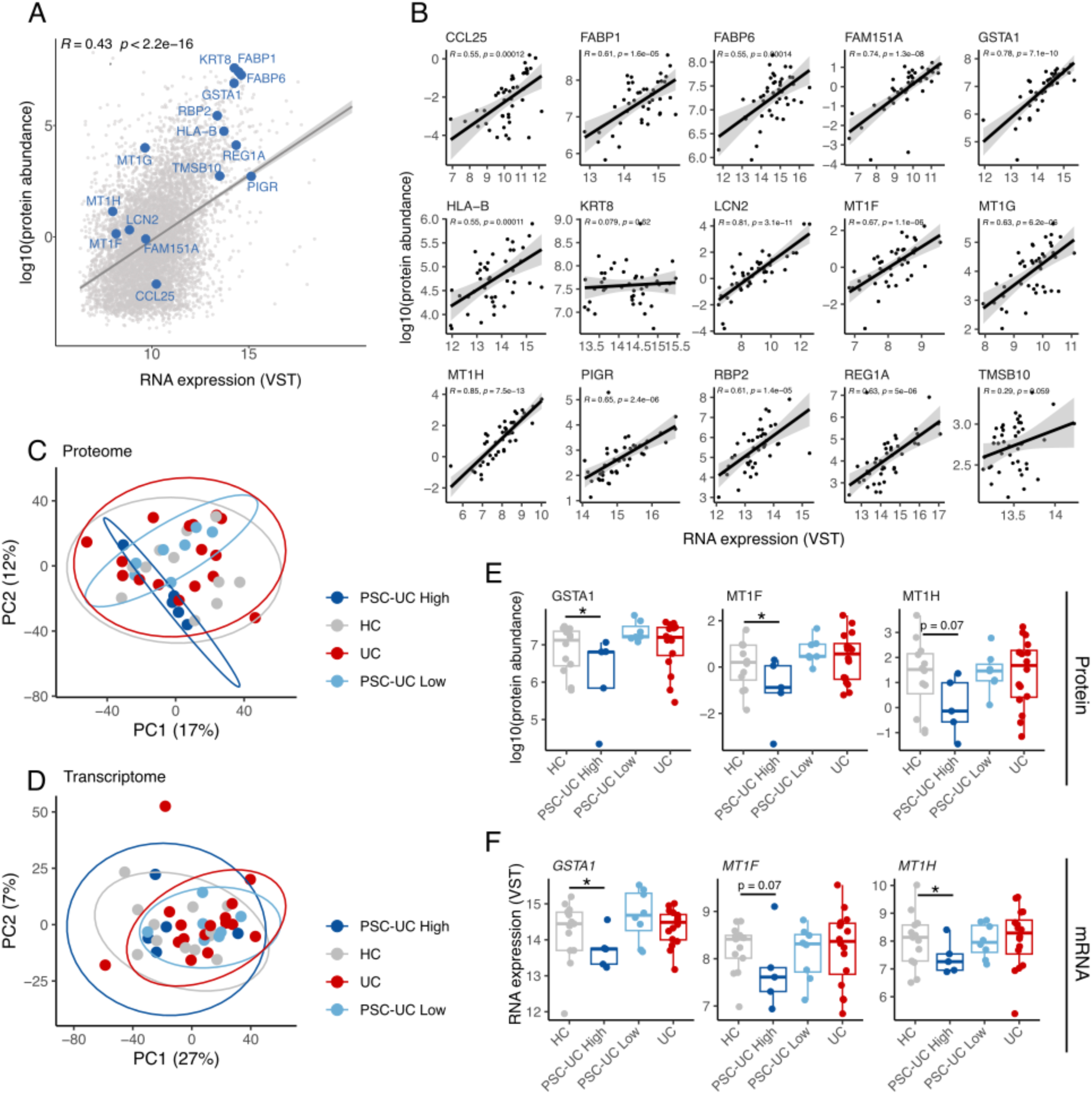
Proteomic analysis supports ALP-associations with detoxification protein abundance. (A) mRNA vs. protein abundance measures for 6080 genes/proteins that were shared between RNA-seq and proteomic datasets. Highlighted are genes that were identified as being co-expressed with *GSTA1* in enterocytes (B) Scatterplots showing the relationship between mRNA and protein abundance across samples. (C) Principal components plot showing data structure according to diagnosis in proteomic data. (D) Principal components plot showing data structure according to diagnosis in transcriptomic data. (E) Abundance of GSTA1, MT1F and MT1H proteins across diagnoses. (F) Abundance of GSTA1, MT1F and MT1H RNA across diagnoses. The PSC-UC group was split into ALP High (>1.5x upper limit of healthy control U/L) and ALP Low (<1.5x upper limit of healthy control U/L) groups. *nominal p < 0.05.

These data support the use of RNA measurements as a proxy for protein abundance for ileal genes associated with detoxification, and suggest that the proteins GSTA1, MT1F and MT1H are reduced in abundance in conjunction with reduced liver function in PSC-UC. This supports a role for reduced bile flow in regulating transcription and abundance of proteins involved in detoxification processes in the human ileum.

### Ileal *Gsta1/2* is decreased in murine cholestatic liver disease

Given our observations of *GSTA1* as a marker of a dominant enterocyte cell state, decreased *GSTA1* mRNA in PSC-UC, and decreased GSTA1 protein with increased ALP, we wanted to further determine if decreased *GSTA1* expression is driven by cholestasis *per se*. To test this, we induced cholestatic liver disease in C57BL/6 mice using a 3,5-Diethoxycarbonyl-1,4-Dihydrocollidine (DDC) diet and compared them to mice on a control diet. At day 5, mice fed DDC displayed had lost significant body weight (**Fig. S6A**) and had histological features of cholestatic liver disease that included cellular proliferation in the portal field and increased collagen deposition (**Fig. S6B**). Consistent with our observations in humans, *Gsta1/2* mRNA levels were significantly lower in mice that had received the DDC-containing diet relative to control animals at day 5 (**Fig. 5A, left panel**), a feature that we also observed in the *Mdr2^−/-^* genetic model of cholestatic liver disease (**Fig. 5A, right panel**). We also observed reduced bile acid transporter, *Slc10a2*, mRNA levels (p = 0.07), but not reduced bile acid binding protein (*Fabp6*) levels in DDC ileal samples (**Fig. S7A-B**).

**Fig. 5.**
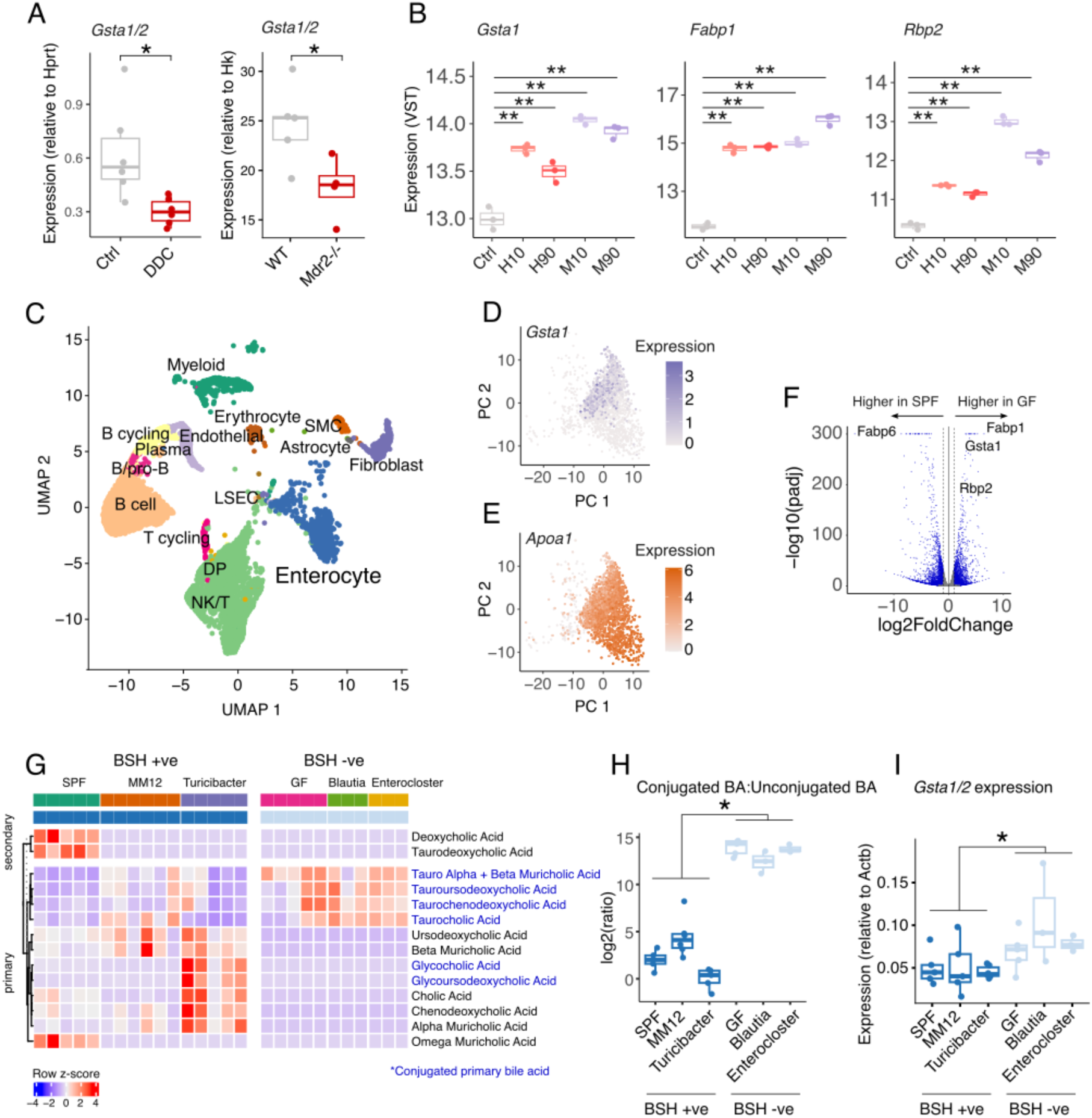
*Gsta1* expression is driven by cholestasis and tuneable by the microbiome. (A) Quantitative PCR analysis of *Gsta1/2* transcription in ileal samples taken from mice fed a control diet or a 0.1% DDC containing diet after 5 days (left panel) and wild-type or *Mdr2*^−\-^ mice (right panel). **p < 0.01 using Wilcox Ranked Sum test. (B) Expression of *Fabp1*, *Gsta1* and *Rbp2* mRNA using publicly available RNA-seq data. Ctrl = micelles without BA, H10 = 10% 12α-hydroxylated BAs in a human bile acid pool, H90 = 90% 12α-hydroxylated BAs in a human bile acid pool, M10 = 10% 12α-hydroxylated BAs in a murine bile acid pool, M90 = 90% 12α-hydroxylated BAs in a murine bile acid pool. **p<0.01 using Wilcoxon Ranked Sum test. (C) Publicly available scRNA-seq data from mouse terminal ileum. Cell annotations from the original publication were used in this analysis. (D) Expression of *Gsta1* in the enterocyte compartment of mouse ileum. (E) Expression of Apoa1 in the enterocyte compartment of mouse ileum. (F) Enterocyte pseudobulk differential expression analysis between GF and SPF. Highlighted in blue are genes that are significantly changed (padj < 0.05 and |log2(fold change)| > 1. (G) Heatmap showing bile acids which are significantly differentially abundant across gnotobiotic mouse models that do (SPF, MM12, *Turicibacter*) and do not (GF, Blautia, Enterocloster) have encoded BSH genes. (H) Ratio of conjugated to unconjugated primary bile acids in different gnotobiotic mouse groups, coloured by BSH status. Significance determined by Wilcoxon Rank Sum Test. (I) qPCR analysis of Gsta1/2 transcription in mouse ileum from gnotobiotic mouse groups. Turicibacter = mice colonised with BSH+ *Turicibacter* UBA1159 (∼87%) and *Mammalicoccus lentus* (∼13%) in their small intestine. MM12 = Oligo Mouse Microbiome 12, Blautia = *Blautia coccoides* (YL58) monocolonised, Enterocloster = *Enterocloster clostridioformis* (YL32) monocolonised. **p < 0.01, *p < 0.05.

### Ileal *Gsta1/2* is regulated by bile acids

We reasoned that reduced expression of ileal *Gsta1* may be due to reduced concentrations of bile acids that are usually transported to the ileum in the absence of cholestasis. To test this, we used publicly available RNA-seq data from murine ileal epithelial organoids stimulated with micelles containing murine (M10 and M90) or human-based (H10 and H90) bile acid pools^34^. We found that stimulation of ileal epithelial organoids with either mouse or human bile acid pools upregulated the expression of *Gsta1* and the *Gsta1*-associated genes, *Fabp1* and *Rbp2,* relative to control samples (micelles lacking bile acids) (**Fig. 5B**), supporting bile acid mediation of transcription. We next wanted to assess the extent to which bile acid modifications - specifically amino acid conjugation - affected the regulation of *Gsta1* and associated genes. To gain a broad understanding, we utilised data from germ-free (GF) mice, which do not have microbiome-dependent bile acid deconjugation capacity (bile salt hydrolases (BSH)). We used publicly available, pre-annotated scRNA-seq data^35^ (**Fig. 5C**), and observed that, similar to humans, murine ileal enterocytes exhibit cell states that are marked by either high *Gsta1* or high *Apoa1* expression (**Fig. 5D**-**E**). Using pseudobulk differential expression analysis, we found that *Gsta1* and the *Gsta1*-associated genes *Fabp1* and *Rbp2* were significantly more highly expressed in GF enterocytes compared to SPF enterocytes (**Fig. 5F**). These data support a regulatory role for conjugated bile acids in mediating *Gsta1* expression. To evaluate this further, we generated gnotobiotic mice with minimal microbiomes with known presence (SPF, Oligo MM12 and *Turicibacter* UBA1159- containing) or absence (*Blautia coccoides* YL58 and *Enterocloster clostridioformis* YL32 monocolonisations) of BSH genes, and assessed ileal expression of *Gsta1/2.* Consistent with Predicted BSH activity in these varied microbiomes, we observed significant increases in conjugated primary bile acids in mice harbouring microbiomes with no BSH encoded genes (**Fig. 5G**). This was confirmed by assessing the ratio of conjugated:unconjugated bile acids in these gnotobiotic mouse models (**Fig. 5H**). Consistent with a role of conjugated primary bile acids on regulating *Gsta1/2* mRNA, mice harbouring microbiomes with known BSH presence had significantly lower expression of *Gsta1/2* than mice lacking BSH (**Fig. 5I**).

Together, these results support reduced bile flow in mediating PSC-associated decreases in *GSTA1*/*Gsta1* mRNA expression, a feature that is likely due to reduced ileal signalling of conjugated primary bile acids.

## Discussion

Development and progression of liver inflammation and damage in PSC and PSC-UC involves pathogenic communication between the gut and the liver which is likely mediated by the gut microbiome. This may be partly driven by inflammatory processes in the colon, where tissue inflammation leads to epithelial barrier dysfunction, increased permeability and systemic exposure to microbial products and/or inflammatory cells. It is noteworthy, however, that increased intestinal permeability is a feature of other chronic liver diseases that do not present with overt intestinal inflammation.^36^ In addition, liver pathology is not a dominant feature of patients with inflammatory bowel disease, suggesting that intestinal inflammation is not sufficient to drive microbe-driven liver inflammation and damage. A recent large intestinal scRNA-seq survey^37^ supports the distinction of the PSC-associated microbiome and mucosal cellular and transcriptomic landscape from UC, which may explain some of the disease-specific tissue involvement.

In the current study, we sought to address whether the terminal ileum plays a somewhat hidden role in PSC. Given previous evidence of small intestinal involvement in chronic liver disease,^38^ an association of ileitis with PSC,^39^ the assessment of ileum-specific and pan-inhibitors of the ileal bile acid transporter, SLC10A2/IBAT, in phase 2 clinical trials^40^ and its critical role in bile acid homeostasis, we wanted to understand any potential role for the terminal ileum in gut-liver communication in PSC.

Using high-throughput assays targeting single cells, spatial niches and protein abundance, we have characterised the terminal ileum in health and PSC-UC. We were able to detect the major absorptive and secretory lineages of ileal epithelial cells in our scRNA-seq atlas and were able to identify previously undefined cell states in ileal enterocytes that were characterised by either high *GSTA1* expression or high *APOA1* expression. Genes that were co-expressed with *GSTA1* included genes for bile acid binding proteins, *FABP1* and *FABP6*, which was the first indication that *GSTA1* and other detoxification enzymes in ileal enterocytes are regulated by bile acids. Reasons for the cell state compartmentalisation of enterocytes that we observed in spatial transcriptomics data are unclear at present, although a previous study in human proximal small intestine reported similar findings, with *GSTA1* being predominantly expressed in the lower portion of villi and *APOA4* being expressed predominantly at the villus tip.^33^ It is intriguing that enterocytes may be differentially sensitive to bile acids along the crypt-villus axis and is a feature of barrier function that should be investigated further.

Differential expression analysis of scRNA-seq data, bulk transcriptomes and proteomes in patient cohorts revealed reduced abundance of detoxification genes/proteins (GSTA1, MT1H, MT1F) in PSC-UC relative to healthy controls, as well as associations with reduced liver function (i.e. high ALP). We do not yet know the consequences of these decreases in detoxification gene expression programmes, although the role of *GSTA1* in reducing reactive oxygen species (ROS) and/or controlling proliferation^41^ could be significant. For example, the ability of enterocytes to control homeostatic production of ROS is integral to maintaining barrier integrity.

We further validated our observations in two mouse models of PSC, demonstrating that cholestasis induced by feeding mice a diet of DDC or genetic deletion of *Mdr2* causes a reduction in *Gsta1/2* expression in the murine ileum. This is likely due to reduced bile acid signalling, as we could demonstrate that bile acid pools are able to drive expression of *Gsta1* and *Gsta1*-associated genes in ileal epithelial organoids. Finally, we have evidence that reduced expression of *Gsta1/2* is related to restricted flow of primary conjugated bile acids from the liver to the intestine in PSC - demonstrated through an increased expression of *Gsta1/2* in the ileum of germ-free mice and BSH-ve gnotobiotic models relative to mice with microbial BSH activity (SPF, MM12 and *Turicibacter* UBA1159-containing minimal microbiome mice).

### Limitations

There are a number of limitations of our study that need to be addressed. Patient cohorts in this study were small, resulting in a limited ability to perform deep and systematic analysis of confounding factors. One such factor is the receipt of ursodeoxycholic acid (URSO) by patients with PSC-UC. This is a secondary bile acid which may have impacts on the ileum that we could not control for in this study and would require follow-up studies with more patients and appropriate controls. We also did not have any patients with PSC only. Samples analysed in the present study were from patients who did not have active IBD, yet we still cannot conclude that PSC-UC with inactive disease is equivalent to PSC without IBD in terms of ileal transcriptomic and proteomic signatures.

*GSTA1* is a very interesting gene which is highly conserved across mammalian species. However, it is part of a large family of glutathione-S-transferases, making homology and human-to-mouse orthology mapping difficult. In our mouse analyses, we used a qPCR probe that recognises exons from both *Gsta1* and *Gsta2*. We therefore do not have a conclusive picture of which homolog is reduced in murine cholestasis.

### Conclusions

Our data support altered barrier function in the ileum of patients with PSC-UC that is linked to epithelial detoxification pathways. Whether these gene expression and protein changes result in decreased barrier integrity that could lead to translocation of bacterial and/or dietary products that impact on liver function and inflammation should be explored in future studies, as well as the potential to enrich for conjugated bile acids through microbial BSH inhibition.^29^

## Materials and methods

Detailed materials and methods can be found in supplementary information.

## Ethics statement

Study participants with PSC-UC, UC or healthy controls were identified via Oxford University Hospitals NHS Trust endoscopy service (Oxford, UK). Intestinal biopsy samples were taken at endoscopy with written informed consent under the Oxford Gastrointestinal Illnesses Biobank study (REC: 21/YH/0206).

## Author contributions

**Miriam Tomczak:** Conceptualisation, Methodology, Formal analysis, Investigation, Writing – Review and Editing. **Katherine Coldwell:** Investigation. **Kyla Dooley:** Investigation, Data Curation. **Amelia Heslington:** Investigation. **Claire Pearson:** Investigation, Resources. **Elizete Araujo:** Investigation. **Megan Perry:** Investigation, Data Curation. **Kaye Eve-Rowe:** Investigation, Data Curation. **Shana-Lee Bownes:** Investigation, Data Curation. **Yukari Kimura:** Investigation, Data Curation, Resources. **Nicholas McDermott Provine:** Investigation. **Lisa Gartner:** Investigation. **Amy Cross:** Investigation. **Joanna Hester:** Investigation. **Iolanda Vendrell:** Investigation, Supervision. **Anna Hoyle:** Investigation, Formal Data Analysis. **Roman Fischer:** Supervision. **Sandi Yen:** Formal Data Analysis. **Charlotte Rich-Griffin:** Methodology. **Ida Parisi:** Investigation, Supervision. **Oyindamola Ajisegiri:** Investigation. **Ethar Ali**: Investigation. **Sumaiya Hoque:** Data Curation. **Fadi Issa:** Supervision. **Samuel Huber:** Supervision. **Fiona Powrie:** Supervision, Resources. **Matthias Friedrich:** Investigation, Supervision, Methodology. **Holm Hans Uhlig:** Supervision. **Calliope Dendrou:** Supervision. **Emma Louise Culver:** Conceptualisation, Supervision, Resources, Project Administration, Funding Acquisition, Writing – Review and Editing. **Paul Klenerman:** Conceptualisation, Supervision, Resources, Project Administration, Funding Acquisition, Writing – Review and Editing. **Nicholas Ilott:** Conceptualisation, Supervision, Resources, Formal Analysis, Project Administration, Funding Acquisition, Writing – Original Draft.

## Data availability

Raw scRNA-seq data are available through ArrayExpress hosted at EMBL-EBI with accession number E-MTAB-17497. Raw bulk RNA-seq data are available through ArrayExpress hosted at EMBL-EBI with accession number E-MTAB-17575. The mass spectrometry proteomics data have been deposited to the ProteomeXchange Consortium via the PRIDE partner repository with the dataset identifier PXD085057. Processed scRNA-seq, bulk RNA-seq and proteomics data are available through figshare (doi: 10.6084/m9.figshare.c.8587205). Code used to analyse the data are available at Github (https://github.com/OxfordCMS/PSC_Ileum_Detox).

## Ackowledgements

We would like to thank the patients who kindly donated samples to the study. We thank Novogene Europe (Cambridge, UK) for the sequencing support. Computational aspects of this work were supported by Oxford’s Biomedical Research Computing (BMRC) facility.

