## Supplementary information for "Bile acid-mediated ileal barrier function deficits in primary sclerosing cholangitis"

### Supplementary methods

**Intestinal biopsy collection, preservation, and dissociation**

Intestinal biopsy samples were collected from the ileum and the caecum during endoscopy and immediately placed in sterile collection medium (RPMI-1640 (Sigma-Aldrich) + 1% human AB serum (Sigma-Aldrich)) on ice for transportation. Samples were cryopreserved in CryoStor Cs10 (STEMCELL Technologies). When required, samples were thawed rapidly in a 37 °C water bath and washed in 10ml R10 culture medium (RPMI-1640 (Sigma-Aldrich) + 10% FBS (Sigma-Aldrich) + 1% penicillin/streptomycin (Sigma-Aldrich)) before tissue dissociation.

For isolation of single cells for single-cell RNA sequencing (scRNAseq), biopsy samples underwent enzymatic and mechanical digestion. Briefly, biopsies were finely minced to sub-mm size using a scalpel and transferred to 1ml R10 medium with 100 µg/ml Liberase TL (Roche) and 100 µg/ml DNase I (Roche). Samples were incubated for 30 minutes at 37 °C on a magnetic stirrer at 1100 rpm, having been passed through an 18G needle at the halfway point. Dissociated cells were passed through a 70-µm filter and washed with ice cold resuspension buffer (PBS (Sigma-Aldrich) + 0.5% BSA (Sigma-Aldrich) + 2mM EDTA (Sigma-Aldrich)) before cell counting using the Countess™ 3 Automated Cell Counter (Invitrogen). Finally, the cells were resuspended in 100-400µl loading buffer (PBS + 0.04% BSA) for scRNAseq.

**Single-cell RNA sequencing**

Up to 20,000 cells per sample were loaded in each channel of the Chromium Next GEM Chip K and the 5' gene expression libraries were generated following the manufacturer's protocol[1] (10X Genomics). Sequencing was carried out by Novogene (Cambridge, UK) on the Illumina NovaSeq 6000 platform, in 150 base pairs paired end dual indexed mode, with read 1 set to

26 cycles, i7 index to 10 cycles, i5 index to 10 cycles and read2 to 90 cycles. With an aim that each cell be represented by at least 20,000 read pairs for the 5' gene expression library, sequencing was carried out to a depth of 120GB for the gene expression libraries. Library loading and pooling was done in accordance with the user guide and a 5% PhiX spike-in was included in each sequencing run.

**scRNA-seq data processing**

Feature-barcode matrices were generated from 10x Genomics Chromium scRNA-seq data using cellranger v7.1.0. We used *cellranger count*, aligning sequenced reads against the GRCh38 genome build, annotated with genes from ENSEMBL 98. To enable manual inspection and filtering of resulting data we employed the *--force-cells=10000* argument to cellranger count to produce feature-barcode matrices with 10,000 cells per sample. Parallelisation of processing was achieved through the implementation of cellranger through the cellhub (https://github.com/sansomlab/cellhub) cgat-core based pipeline. We initially defined and removed doublets using *scrublet*[1] v0.2.3 with an expected doublet rate of 0.06, and min_counts=2, min_cells=3, min_gene_variability_pctl=85 and n_prin_comps=30 passed to the *scrub_doublets* function. We used Seurat[2] v5.2.1 in R 4.3.1 to asses cell quality in terms of proportion of mitochondrial reads and RNA counts. Given the complexity of cell types in biopsy samples we initially used permissive mitochondrial read filtering (<75%) in combination with RNA counts per cell required to be between 500 and 5000. This left a total of 17,329 (median = 1,387/patient) cells for the ileum and 11,869 (median = 717/patient) for the caecum.

**Clustering and sub-clustering of scRNA-seq data**

Cells derived from ileal and caecal samples were clustered independently of each other using a Seurat-based workflow that included harmonising data to account for individual differences. Data were log normalised, scaled and variable features were determined using the *FindVariableFeatures* function, specifying 2000 variable features. Dimensionality reduction was performed using the *RunPCA* function and data were harmonised on patient identifier using Harmony[3] v1.2.0 (*RunHarmony* function in Seurat). To identify initial cell clusters we used the *FindNeighbours* function in Seurat using the harmony reduction and the first 30 dimensions. Following this, we identified clusters using *FindClusters* with Leiden clustering at a resolution of 0.5. To annotate the resulting clusters, we used cluster markers found by *FindAllMarkers* in combination with the literature[4,5]. We first broadly annotated the clusters using markers for lymphocytes, myeloid cells, epithelial cells and stromal cells before sub-clustering each compartment. After broad annotation, we split the data into immune, epithelial and stromal compartments for sub-clustering. For the immune compartment, we selected cells that were annotated as B cells, T cells, monocytes/macrophages or mast cells. We removed cells with >10% mitochondrial reads, re-clustered them (2000 variable features, harmony integration using 30 PCA-derived dimensions and Leiden clustering at a 0.6 resolution). We annotated the resulting clusters using marker genes from *FindAllMarkers* and from the literature. For epithelial cells, we did not lower the proportion mitochondrial read threshold for sub-clustering and proceeded to cluster re-cluster and annotate cells as we did for the immune cell compartment.

**Differential composition analysis**

To assess differences in cell frequencies between disease states we used propeller[6], performing analysis on immune cells, epithelial cells and stroma/endothelial cells in separate analyses. Differences in cell proportion were determined at a significance of $p < 0.05$.

#### Enterocyte cell state analysis

To explore genes associated with *GSTA1* and *APOA1* enterocyte cells we used scWGCNA v1.0.0 (https://github.com/CFeregrino/scWGCNA). This is an adaptation of Weighted Gene Co-expression Network Analysis (WGCNA), originally designed for bulk RNA-seq and microarray data[7]. We downloaded a scRNA-seq dataset that was specifically designed to look at human epithelial cells, GSE125970[4] to maximise power to observe and analyse different cell states. We ran scWGCNA in R 4.3.1 using the *run.scWGCNA* function which performed the correlation analysis on normalised data. We searched the resulting co-expression modules for *GSTA1* and *APOA1* and considered genes associated with these two cluster markers if they had a module membership (i.e. correlation with module eigengene) of >0.6. Confirmation of co-expression was performed in our data through visualisation in heatmaps. Functional enrichments in modules of genes co-expressed with *GSTA1* or *APOA1* were assessed using overrepresentation analysis (implemented using the *fora* function in fgsea[8]) of these gene sets against Gene Ontology (GO) biological pathways (BP) obtained from the MSigDB (https://www.gsea-msigdb.org/gsea/msigdb, v.7.1, c5.bp). Pathways were considered significantly overrepresented at an adjusted p-value < 0.05.

#### Pseudobulk differential expression analysis

For each cell compartment described in the results section we performed pseudobulk differential expression using samples that had at least 20 cells for that compartment. Counts were aggregated at the patient-level using the *AggregateExpression* function in Seurat. Genes were kept for differential expression analysis if they had at least 10 counts in at least the minimum number of samples for a given group (n = 2-4 depending on cell type). We used DESeq[9] v1.42.1 in R 4.3.1 for differential expression analysis, using the *lfcShrink* function to obtain moderated fold change estimates for each gene. Genes were considered differentially expressed at a DESeq2 adjusted p-value < 0.05. To determine pathway enrichment, we used

Gene Set Enrichment Analysis (GSEA) implemented in the R package fgsea[8]. We used log2(fold change) of PSC-UC vs. Healthy controls as the value for ranking and assessed significance of GO BP gene sets obtained from MSigDB v7.1 (https://www.gsea-msigdb.org/gsea/msigdb, c5.bp). Pathways were considered significant at an adjusted p-value < 0.05.

**Sample preparation for Nanostring GeoMX spatial transcriptomics analysis**

Ileal biopsies were collected and immediately placed in 1% formalin. Samples were transferred into 70% EtOH after 24 hours of fixation in formalin andd embedded in paraffin blocks. 5 µm sections were cut and placed on Leica adhesive microscopic slides and baked overnight at 60 °C. Slide preparation was conducted according to NanoString's protocol. Slides were deparaffinized and rehydrated. Target retrieval was performed using IHC Antigen Retrieval Solution (eBioscience) for 20 min (100 °C), followed by 15 min (37 °C) in 1 µg $ml^{-1}$ Ambion Proteinase K (ThermoFisher Scientific). After retrieval, slides were fixed in 10% neutral buffered formalin and washed. Samples were UV light-treated (405 nm, 24 h) to quench background autofluorescence. Next, slides were incubated with human Whole Transcriptome Atlas probes (NanoString) for 16 h (37 °C).

Slides were washed in formamide-SCC buffer before tissue blocking and immunofluorescent staining in Buffer W (Nanostring) with 1% Fc-Receptor block (Miltenyi) and 5% donkey serum (blocking buffer). The sections were then incubated with 1:200 anti-CD68 (SantaCruz, mouse, [KP1]) and 1:200 anti-CD3 (Abcam, rabbit, [SP162]) for 1 h; followed by 1:1000 anti-mouse-AF647, 1:1000 anti-rabbit-Cy3, 1:40 CD45-AF594 (Nanostring) and 1:20,000 Sytox Green for 1 h. All antibody incubations were done in blocking buffer at RT. Epithelial cells were stained by: 1:200 anti-panCK (AE-1/AE-3)-DyLight594 (mouse, Novus NBP2-332000).

Slides were imaged with the Nanosting GeoMx Digital Spatial Profiler with manual selection of regions of interest which included regions that represented villi tips and bases that were marked by high PanCK signal. Segmentation of these regions allowed for the acquisition of oligonucleotide probes from PanCK+ cells which we used for downstream analysis. For library generation, samples were subjected to PCR using i5 and i7 dual indexing primers (Nanostring) before pooling and purification using AMPure XP beads (Beckman Coulter). Library QC was done using Qubit (ThermoFisher Scientific) and TapeStation (Agilent). Resulting libraries were sequenced on the Illumina NovaSeq platform using 150-bp paired-end sequencing.

**Nanostring GeoMX data processing**

Fastq files per segment were processed into Digital Count Conversion (DCC) files using the Nanostring GeoMxNGSPipeline tool v2.3.3.10. After processing, the counts data were integrated with probe annotations and segment metadata by creating a NanoStringGeoMxSet R object (using the Hs_R_NGS_WTA_v1.0.pkc file to map probes to target genes). Quality control and pre-processing of the data were performed using Shiny geomx (https://github.com/OxfordCMS/OCMS_Shiny_Geomx), an R/Shiny wrapper and extension for the GeomxTools R package. Default values were used for filtering samples and probes with the exception that probes had to be expressed at least 1 standard deviation above negative control probes to be taken forward for downstream analysis. Quantile normalisation was used to account for between-segment variability. To assess for differences in expression between PanCK+ cells in villus tips vs. bases, we subset the data for genes that were present in either the *GSTA1* or *APOA1*-associated module and performed differential expression using linear mixed-effects models with slide.name as a random effect implemented using the lmerTest package in R 3.2.1.

#### Analysis of publicly available human ileum Visium data

Ileum data were downloaded from https://data.mendeley.com/datasets/v8s9nz948s/1. This dataset consists of Visium spatial transcriptomic data on serial sections from the same ileal sample. To visualise the spatial distribution of *APOA1* and *GSTA1*, we loaded the data into Seurat v5.6.0, R4.5.1. Location of expression relative to villi was visualised using the *SpatialFeaturePlot()* function. Presented are the data from the section with identifier V19T26-028_B1.

#### Samples for bulk proteomics and RNA-seq

Ileal biopsies were collected from patients undergoing routine colonoscopy (n = 14 PSC-UC, n = 17 UC and n = 14 HC). Samples were stored in RNALater at 4°C before being transferred to -80°C before RNA and protein extraction.

#### RNA and protein extraction

Tissue samples were thawed on ice and RNA was extracted using the Qiagen RNeasy mini kits according to the manufacturer's instructions. The only exception to this was that we collected the first flow-through from the column which contained proteins. These protein-containing samples were maintained at -80°C until protein extraction and liquid chromatography coupled with mass-spectrometry (LC-MS) analysis. After elution, RNA was stored at -80°C until sequencing. RNA samples were sent to Novogene (Cambridge, UK) for mRNA sequencing. Briefly, Messenger RNA was purified from total RNA using poly-T oligo-attached magnetic beads. After fragmentation, the first strand cDNA was synthesized using random hexamer primers followed by the second strand cDNA synthesis. The library was ready after end repair, A-tailing, adapter ligation, size selection, amplification, and purification. The library was checked with Qubit and real-time PCR for quantification and bioanalyzer for size distribution detection. Quantified libraries were pooled and sequenced on Illumina Novaseq Plus at a length of 150bp paired-end reads.

**Bulk RNA-seq data analysis**

We assessed the quality of raw data using Fastqc (https://www.bioinformatics.babraham.ac.uk/projects/fastqc/) v0.11.9. All samples were of high quality and reads reached Q>30 across the length of the reads. We trimmed Illumina universal adapters using trimgalore v0.6.10 with the --length 75 --illumina parameters set. We performed pseudoalignment of sequences to against the human transcriptome (hg38, ENSEMBL 106) using kallisto[10] and the in stranded mode (--rf-stranded). Differential expression analysis was performed using DESeq2 using count data imported using the tximport R package[11]. Age and sex were included in the design as covariates when assessing group differences (design = ~age + sex + diagnosis).

**Proteomics sample preparation**

Protein was precipitated in 4 volumes of ice-cold acetone and the pellet was then spun down and dried. 6M urea in 100mM TEAB was used to resuspend the pellet followed by reduction and alkylation with 0.5M TCEP and Iodoacetamide respectively. Samples were then diluted in 50mM TEAB and digested overnight using trypsin at a ratio of 1:25. This reaction was quenched with 1% TFA and samples were desalted using SOLA SPE cartridges, dried down and resuspended in 0.1% formic acid.

**Proteomics data analysis**

Raw data was searched using DIA-NN (v1.8.1) against UniProt/SwissProt human database (2024) with added decoys and known contaminants which contained a total of 40936 sequences (50% decoys). Searches used a maximum false discovery rate of 1%. Carbamidomethyl was added as a fixed modification to cysteine residues (+57.021Da). A fully tryptic search was conducted with 2 missed cleavages allowed. Data was exported and onward analysis was conducted in Perseus (v2.0.11) where data was normalised by median subtraction and data was imputed based on standard distribution for proteins detected in at least 7 samples. To minimise variability across samples, resulting data were quantile normalised. Differential protein abundance was assessed using linear models implemented using the *lm* function in R 4.3.1. Age and sex were accounted for in the models (formula = ~ age + sex + diagnosis).

**Mice**

Mouse work was conducted at The Kennedy Institute of Rheumatology (KIR) at the University of Oxford or at the animal facility of the University Hospital Hamburg-Eppendorf. Experiments were conducted in accordance with local animal care committees (UK Animal Scientific Procedures Act of 1986). Mice were routinely screened for the absence of pathogens, and were kept in individually ventilated cages with environmental enrichment. Female, 8-12 week old, C57BL/6J specific pathogen free (SPF) mice were bred in-house and provided with sterile water and food irradiated to 50kGy *ad libitum*. A group of mice were fed 3,5-diethoxycarbonyl-1,4-dihydrocollidine (DDC)-containing diet irradiated to 50kGy. At the time of humane killing, mice were killed by asphyxiation using a rising concentration of carbon dioxide, authorised via Schedule 1 of A(SP)A 1986, and death confirmed via cervical dislocation.

**Experimental model of cholestatic liver disease**

N = 5 mice (Ctrl) were fed standard diet for 5 days and compared to n = 7 mice that received DDC over the same time period. Weight change was recorded daily and mice were killed if they fell below 85% of their starting weight. All mice were killed 5 days after starting the experiment. Liver pathology in the two groups was assessed by haematoxylin and eosin

(H&E) and Sirius Red staining of liver sections that had been fixed in 1% formalin, transferred to 70% EtOH and paraffin embedded (performed by the histology core facility at the Kennedy Institute of Rheumatology). Evidence of cell expansion, thickening of vessel walls and deposition of collagen was considered as evidence of liver pathology.

**_Mdr2_ $^{-/-}$ genetic model of cholestatic liver disease**

C57BL/6 MDR2 knockout (*Mdr2* -/-) mice were bred and housed under specific pathogen-free conditions (SPF) at the animal facility of the University Hospital Hamburg-Eppendorf. After weaning (at 3-4 weeks of age), mice were separated with respect to their genotype. Mice were sacrificed at 12 weeks of age. Age and sex-matched littermates were used. Food and water were provided *ad libitum*.

**Gnotobiotic mouse models**

Mouse experiments were performed with 6-12 week old female wild-type C57BL/6J mice that were bred and maintained in germ-free isolators, Mice were colonised with specific gnotobiotic strains or consortia through vertical transmission from parents. Mice were colonised with oligo-MM12, a defined consortium of 12 species of bacteria[12], or monocolonised with *Blautia (pseudo)coccoides* YL58, *Enterocloster clostridioformes* YL32 or *Turicibacter* UBA1159 (~87% relative abundance in small intestine contents) and *Mammalicoccus lentus* (~13% relative abundance in small intestine contents). Mice were housed in sterile, flexible film isolators (NKP isotec) with enrichment, irradiated food and autoclaved water. Germ free animals were regularly tested for contamination through aerobic and anaerobic culturing and 16S qPCR, both in-house and outsourced (Fera). Colonised animals were regularly tested through shallow shotgun sequencing (Transnetyx) for the presence of microbes.

**Analysis of small intestinal bile acids in gnotobiotic mouse models**

After humane killing, small intestines were removed and small intestine contents were collected through manual squeezing along the length of the small intestine. Contents were collected into sterile eppendorf tubes and frozen on dry ice. Samples were placed at -70°C until processing. Bile acid profiling was performed by Metabolon (Morrisville, USA) using the Metabolon LC-MS/MS method TAM178: “LC-MS/MS Method for the Quantitation of Bile acids”. This method quantifies primary and secondary bile acids and their conjugates: cholic acid (CA), chenodeoxycholic acid (CDCA), deoxycholic acid (DCA), lithocholic acid (LCA), ursodeoxycholic acid (UDCA), glycocholic acid (GCA), glycochenodeoxycholic acid (GCDCA), glycodeoxycholic acid (GDCA), glycoursodeoxycholic acid (GUDCA), taurocholic acid (TCA), taurochenodeoxycholic acid (TCDCA), taurodeoxycholic acid (TDCA), taurolithocholic acid (TLCA), tauroursodeoxycholic acid (TUDCA), glycolithocholic acid (GLCA), α-muricholic acid (AMCA), β-muricholic acid (BMCA), ω-muricholic acid (OMCA), tauro-α-muricholic acid (TAMCA), and tauro-β-muricholic acid (TBMCA). TAMCA and TBBCA were quantified together as they could not be separated chromatographically. Extrapolated data were used for downstream analysis.

**Statistical analysis of bile acid profiles**

Bile acids were retained if they were above 10ng/mg concentration in at least 3 samples (the minimum number of samples for a group). Significantly differentially abundant bile acids were determined using a Wilcoxon Rank Sum Test implemented in R4.3.1 across all groups: SPF, oligo-MM12, *Turicibacter*, *Blautia (pseudo)coccoides* YL58, *Enterocloster clostridioformes* YL32 and GF as described previously. Bile acids were deemed significantly different at a Benjamini-Hochberg adjusted p-value < 0.05. To determine the extent to which we controlled for deconjugation activity of the microbiomes, we summed the total conjugated and the total unconjugated primary bile acids and created a ratio of conjugated:unconjugated primary bile acids. A Wilcoxon Rank Sum Test (R4.3.1) was used to determine a significant difference in this ratio between microbiomes that do and do not have predicted bile salt hydrolase activity.

##### qPCR analysis of *Gsta1/2* in murine ileum

For all qPCR experiments, RNA was extracted from ileal tissue using QIAGEN RNeasy mini kits (QIAGEN, Germany) according to the manufacturer's instructions. cDNA was generated using the High-Capacity cDNA Reverse Transcription Kit (Applied Biosystems, Thermofisher, ref: 4368813), following the manufacturer's instructions.

For quantitative real-time PCR analysis, TaqMan Fast Advanced Master Mix (Thermofisher) was used. We used the TaqMan probe Mm03646559_mH to approximate *Gsta1* mRNA expression in the ileum. Relative expression was normalized to Hprt (TaqMan probe Mm03024075_m1) and calculated using the $2^{-DCt}$ method. Given the sequence similarity between *Gsta* genes, this probe also targets *Gsta2,* and as such, we refer to the results as *Gsta1/2*.

##### Publicly available data on bile-acid induced gene expression in murine ileal organoids

We downloaded data with accession GSE144398 from the Gene Expression Omnibus (GEO). Pre-processed data (counts) were used for downstream analysis. We transformed the data using the variance stabilising transformation (VST) implemented in DESeq2. To determine whether bile acids induced genes of interest in ileal organoids, we used Wilcoxon Rank Sum tests for *Gsta1*, *Rbp2* and *Fabp1* comparing each BA pool against control samples (micelles not containing BAs). Details of the composition of BA pools can be found in the original publication[13].

##### Publicly available ileal scRNA-seq data in SPF and GF mice

To test whether *Gsta1* and *Gsta1*-associated genes are mediated by the microbiome we downloaded publicly available scRNA-seq data (https://ftp.cngb.org/pub/CNSA/data5/CNP0005580/Single_Cell/CSE0000406/) that contained data from ileal samples[14]. We used existing cell annotations in the dataset. To compare GF to SPF mice, we subset the data for enterocytes and aggregated counts per sample using the *AggregateExpression* function in Seurat. We then performed differential expression analysis between GF and SPF mice using DESeq2. Genes were considered differentially expressed at an adjusted $p < 0.05$.

### Supplementary Figures

#### Fig. S1

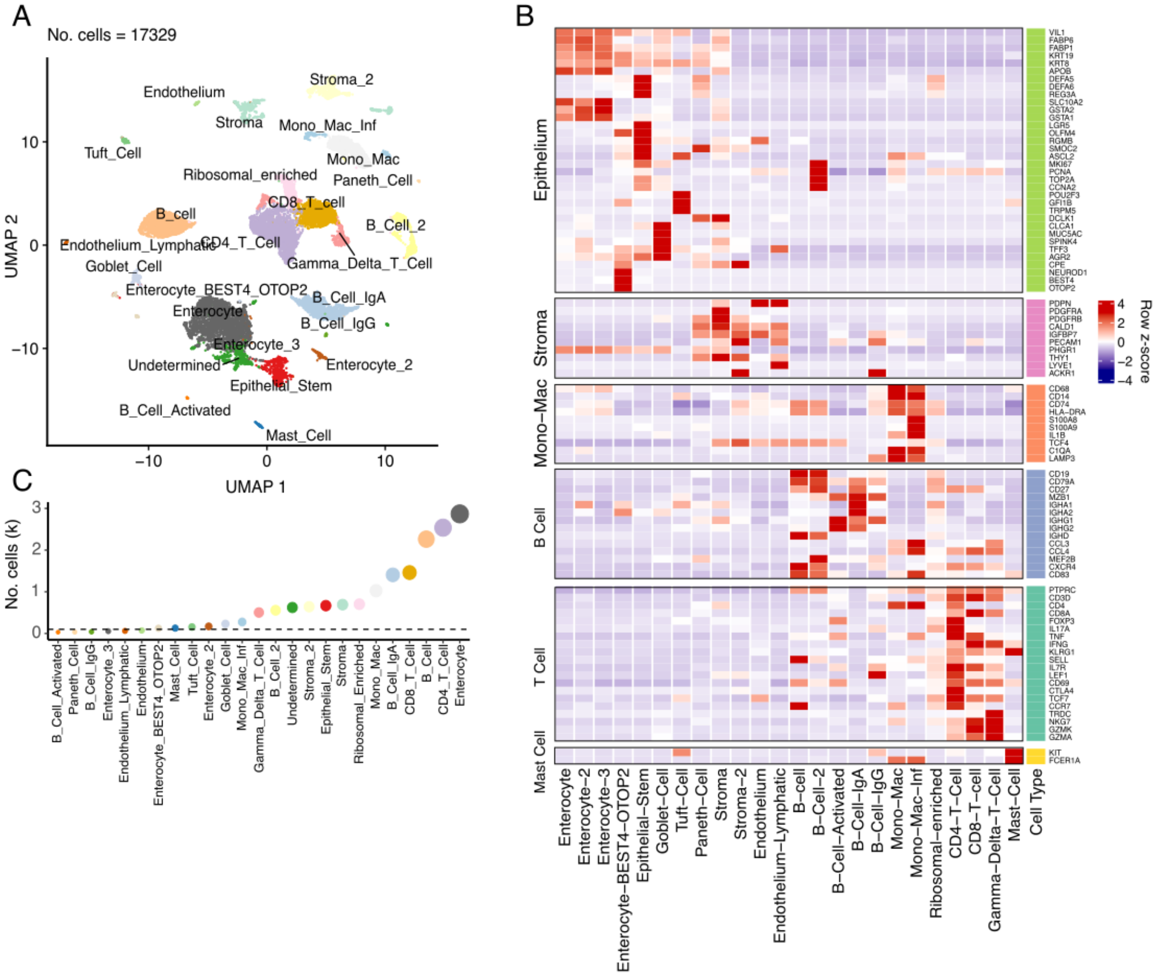


**Fig. S1. A single cell atlas of the terminal ileum in PSC-UC, UC and health.** (A) Broad annotation of cells generated through 10X chromium gene expression profiling. Permissive filtering parameters implemented in Seurat (v5) were applied (%Mt < 75%, No. features > 500, No. UMIs < 5000) to maintain diverse cell types in the analysis. (B) Genes derived from the literature and *de novo* cluster markers used to annotate clusters. (C) Total number of cells (across all samples) that were assigned to each annotated cluster. After initial annotations, cells were subset into their broad cell types and sub-clustered using more stringent and cell-type specific filtering and clustering parameters.

**Fig. S2**

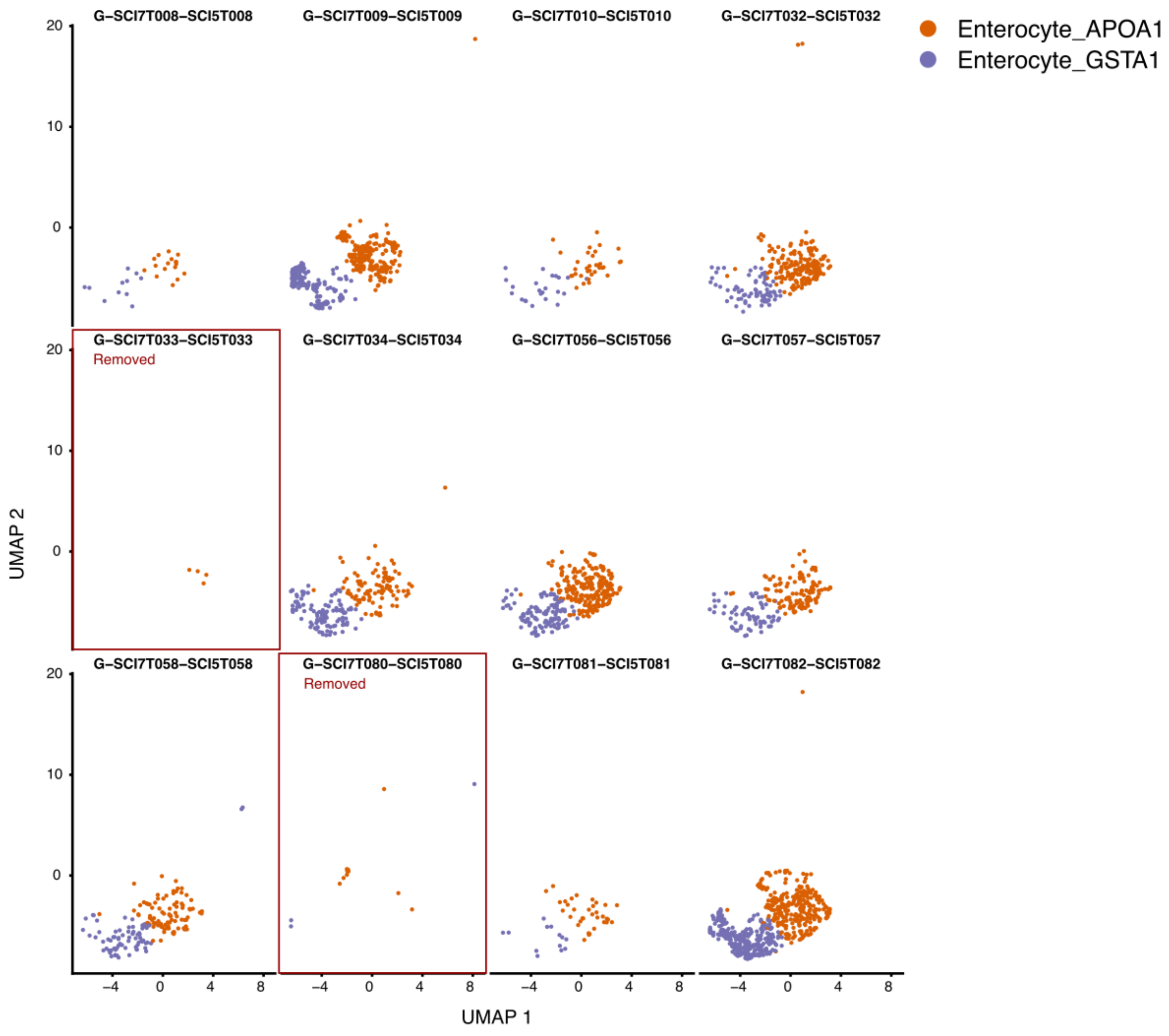


**Fig. S2. Sample removal based on lack of enterocyte states of interest.** Plots represent per sample UMAPs with cells that were annotated as wither *APOA1* high enterocytes or *GSTA1* high enterocytes. Two samples were removed based on their different overall epithelial profiles and lack of representative ileal enterocyte cell states (n = 1 PSC-UC and n = 1 UC).

### Fig. S3

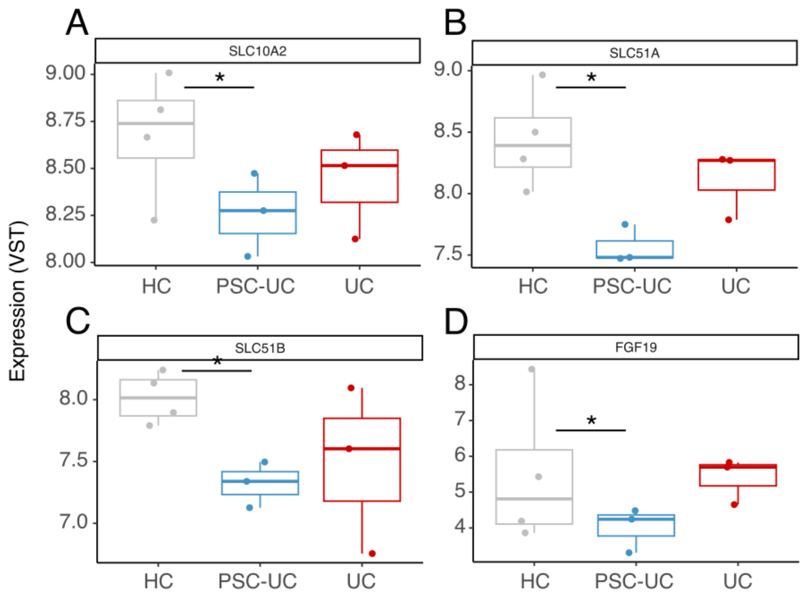


**Fig. S3. Expression of bile acid related genes are decreased in PSC-UC.** Aggregated expression across enterocytes for each sample shows expression differences in (A) *SLC10A2* (B) *SLC51A* (C) *SLC51B* and (D) *FGF19*. *nominal $p < 0.05$.

**Fig. S4**

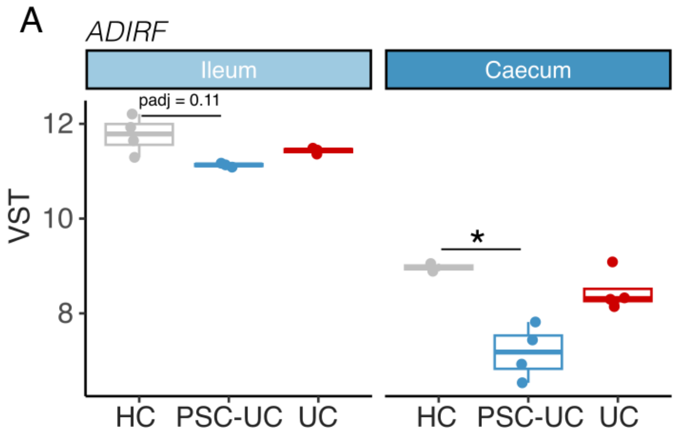


**Fig. S4. *ADIRF* is differentially expressed in absorptive epithelial cells in the ileum and caecum.** (A) Aggregated expression across enterocytes and differential expression analysis for *ADIRF* across ileum and caecum samples. *nominal $p < 0.05$.

**Fig. S5**

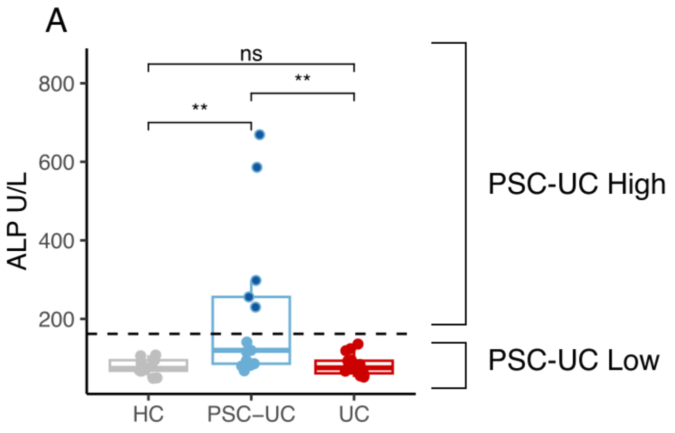


**Fig. S5. Serum alkaline phosphatase (ALP) varies in PSC-UC.** (A) Clinical measurements of serum ALP in participants in the study. The dashed line represents 1.5x the upper limit of healthy control samples.

#### Fig. S6

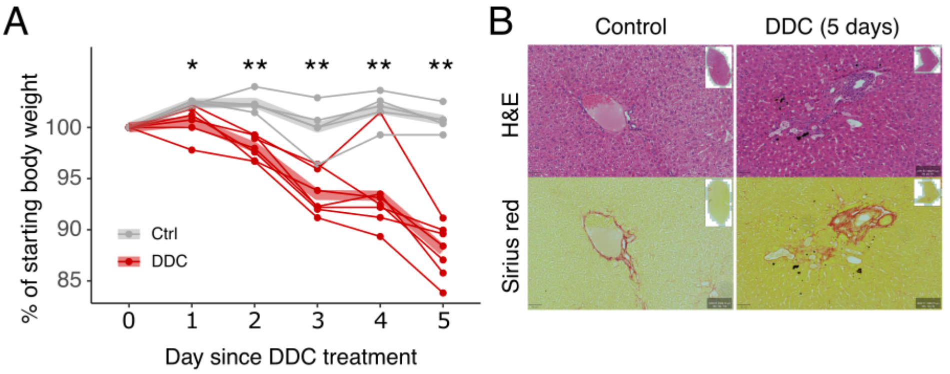


**Fig. S6. DDC model of cholestatic liver disease.** (A) Proportion of starting body weight over time in C57BL/6 mice fed either a control diet or 0.1% DDC containing diet (n = 5-7/group). (B) Representative H&E and Sirius red staining of liver sections from mice fed a standard control diet or 0.1% DDC containing diet after 5 days.

#### Fig. S7

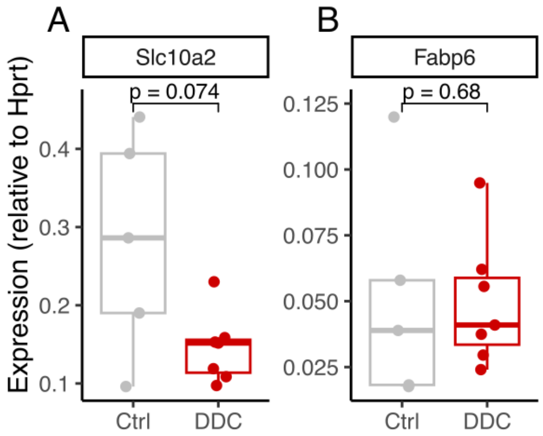


**Fig. S7. Differential expression of bile acid related genes in murine cholestasis.** qPCR analysis of the bile acid transporter (A) *Slc10a2* and bile acid binding protein (B) *Fabp6*.
